# Neuronal and behavioral responses to naturalistic texture images in macaque monkeys

**DOI:** 10.1101/2024.02.22.581645

**Authors:** Corey M. Ziemba, Robbe L. T. Goris, Gabriel M. Stine, Richard K. Perez, Eero P. Simoncelli, J. Anthony Movshon

**Author notes:** These authors contributed equally (Corey M. Ziemba, Robbe L. T. Goris).

## Abstract

The visual world is richly adorned with texture, which can serve to delineate important elements of natural scenes. In anesthetized macaque monkeys, selectivity for the statistical features of natural texture is weak in V1, but substantial in V2, suggesting that neuronal activity in V2 might directly support texture perception. To test this, we investigated the relation between single cell activity in macaque V1 and V2 and simultaneously measured behavioral judgments of texture. We generated stimuli along a continuum between naturalistic texture and phase-randomized noise and trained two macaque monkeys to judge whether a sample texture more closely resembled one or the other extreme. Analysis of responses revealed that individual V1 and V2 neurons carried much less information about texture naturalness than behavioral reports. However, the sensitivity of V2 neurons, especially those preferring naturalistic textures, was significantly closer to that of behavior compared with V1. The firing of both V1 and V2 neurons predicted perceptual choices in response to repeated presentations of the same ambiguous stimulus in one monkey, despite low individual neural sensitivity. However, neither population predicted choice in the second monkey. We conclude that neural responses supporting texture perception likely continue to develop downstream of V2. Further, combined with neural data recorded while the same two monkeys performed an orientation discrimination task, our results demonstrate that choice-correlated neural activity in early sensory cortex is unstable across observers and tasks, untethered from neuronal sensitivity, and thus unlikely to reflect a critical aspect of the formation of perceptual decisions.

**Significance statement:** As visual signals propagate along the cortical hierarchy, they encode increasingly complex aspects of the sensory environment and likely have a more direct relationship with perceptual experience. We replicate and extend previous results from anesthetized monkeys differentiating the selectivity of neurons along the first step in cortical vision from area V1 to V2. However, our results further complicate efforts to establish neural signatures that reveal the relationship between perception and the neuronal activity of sensory populations. We find that choice-correlated activity in V1 and V2 is unstable across different observers and tasks, and also untethered from neuronal sensitivity and other features of nonsensory response modulation.

## Introduction

As signals propagate along the visual hierarchy, individual neurons represent increasingly complex aspects of the visual environment. In principle, these later representations are better suited to support complex perceptual tasks than the simpler representations that precede them, and may thus have a more direct relationship with perceptual experience. This relationship is best assessed through the simultaneous recording of neural responses and behavioral reports of perceptual decisions (Newsome et al., 1989; Parker and Newsome, 1998). In many areas of the brain, the activity of individual neurons approaches or even exceeds the perceptual sensitivity of the subject and, moreover, predicts behavioral choices even on trials where the visual stimulus is ambiguous (Britten et al., 1992; Prince et al., 2000; Uka and DeAngelis, 2006; Nienborg and Cumming, 2006, 2014). Such results have been taken as evidence of an area’s participation in the formation of a perceptual decision, regardless of the exact origin of choice-correlated activity Cumming and Nienborg (2016). Whether sensory noise is fed forward to causally influence downstream areas that integrate evidence into a decision (Shadlen et al., 1996), or whether choice-related activity is fed back from decision circuits to sensory areas to support hierarchical probabilistic inference (Nienborg and Cumming, 2009; Haefner et al., 2016), most explanations of choice-related activity suggest it is a reflection of the decision-making process.

A particularly successful application of this approach has been in the investigation of the neural representation of binocular disparity. Both the primary (V1) and secondary (V2) visual cortex are sensitive to visual disparities, but there is a clear shift from absolute to relative disparity sensitivity in V2 compared with V1, more closely aligning with perception (Thomas et al., 2002). Nienborg and Cumming (2006) further demonstrated that V2 neurons are more sensitive to disparity than V1 neurons, and predict behavioral choices in response to ambiguous stimuli, while V1 neurons do not. We wondered whether a similar V1-V2 distinction might exist in the representation of visual form. We previously found that V1 and V2 neurons can be distinguished based on their responses to naturalistic texture stimuli nearly as well as based on their sensitivity to relative disparity (Freeman et al., 2013). Further, multiple observations provide indirect evidence for a relationship between the response of populations of V2 neurons and the perception of naturalistic textures (Freeman and Simoncelli, 2011; Freeman et al., 2013; Ziemba et al., 2016; Ziemba and Simoncelli, 2021). Despite being recorded under anesthesia, V2 responses, but not V1 responses, predicted human psychophysical performance on a naturalistic texture discrimination task (Freeman et al., 2013). Here, we directly test the strength of this link between V2 responses and the perception of naturalistic image structure by measuring neuronal and behavioral sensitivity simultaneously in the same observers.

We found that V2 neurons were substantially more sensitive to naturalistic image structure than V1 neurons, confirming our previous results in anesthetized animals (Freeman et al., 2013; Ziemba et al., 2018, 2019). However, average sensitivity in both V1 and V2 was far from behavior. Correspondingly, we found inconsistent evidence for a relationship between neuronal responses and behavioral choice in either V1 or V2. When compared with previous data collected from the same two monkeys performing an orientation discrimination task, we found that choice-correlated neural activity was unstable across tasks and observers, and untethered from both neuronal sensitivity and signatures of nonsensory modulation. Our results provide further evidence for the functional differentiation of V1 and V2 in the neural representation of naturalistic image structure, but suggest that further elaboration of these signals in downstream areas may more directly support perception of these visual features (Okazawa et al., 2015, 2017; Movshon and Simoncelli, 2015). Finally, we conclude more broadly that the presence of choice-correlated activity in the sensory cortex is unlikely to be a universal indicator of a neural population’s participation in the formation of a perceptual decision.

## Materials and Methods

### Physiology

Two male macaque monkeys (one *M. mulatta* and one *M. nemestrina*) were used in this study. Both animals were previously trained to perform an orientation discrimination task, and neuronal responses recorded from V1 and V2 during task behavior have been previously published (Goris et al., 2017). Here, both animals were retrained to perform a naturalistic texture discrimination task. Experimental procedures conformed to the National Institutes of Health *Guide for the care and use of laboratory animals* and were approved by the New York University Animal Welfare Committee. Under general anesthesia, the animals were implanted with a titanium head post and recording chamber (Adams et al., 2007, 2011). Extracellular recordings were made with dura-penetrating glass-coated tungsten microelectrodes (Alpha Omega), advanced mechanically into the brain. We distinguished V1 from V2 on the basis of depth from the cortical surface and changes in the receptive field location of the recorded units. We made recordings from every unit with a spike waveform that rose sufficiently above noise to be isolated, and only included recordings in the final analysis that maintained sufficient isolation throughout the session. While the monkey fixated a central fixation spot, we first used a drifting sinusoidal grating stimulus to map each isolated unit’s receptive field and determine where to position the stimulus center in the main experiment. Across our population, receptive fields were centered at eccentricities ranging from 2° to 6°from the center of gaze. Thereafter, we ran an initial characterization of the sensitivity of each neuron to naturalistic image structure.

### Stimulus synthesis, presentation, and tuning

We generated naturalistic texture images using the procedure developed by Portilla and Simoncelli (2000). We generated spectrally matched noise stimuli, and stimuli lying between these and the naturalistic textures, as described by Freeman et al. (2013). For each texture naturalness level, we generated synthetic images at a size of 512 x 2048 pixels, and displayed different samples of texture by changing the center of a 4° raised cosine aperture. To determine the tuning of each recorded neuron, we presented a random sequence of different samples of naturalistic and spectrally matched noise stimuli from five different families. We used the five texture families that demonstrated the strongest average naturalistic texture modulation in area V2 in a previous study in anesthetized monkeys (Freeman et al., 2013). Each image was presented for 100 ms with 100 ms of intervening mean luminance. We computed the discriminability between the average firing rate across samples of naturalistic stimuli and spectrally matched noise for each of the five families. We then selected the family for which naturalistic and spectrally matched noise were most discriminable for use in the texture discrimination task.

### Behavioral task

Subjects were seated in a darkened room in front of a gamma-corrected CRT monitor (iiyama HM204DTA) with their heads stabilized. Eye position was recorded with a high-speed, high-precision eye tracking system (EyeLink 1000). We presented visual stimuli at a viewing distance of 57 cm, a spatial resolution of 1,280 *×* 960 pixels, and a refresh rate of 120 Hz. Stimuli were presented using Expo software (http://corevision.cns.nyu.edu/expo/) on an Apple Macintosh computer.

Each trial in the discrimination task began when subjects fixated a small white point (0.2° diameter) at the center of the screen (Fig. 2a). After 250 ms, two choice targets appeared, one on each side of the fixation point (on the horizontal meridian, at 3.5° eccentricity). The choice targets were samples of naturalistic and spectrally matched noise presented in a 2° aperture. The particular samples and the assignment of choice targets to the left or right of fixation were fixed within a session but randomly varied across sessions. After a 500 ms delay, the target appeared. The target was a sample of texture of intermediate naturalness presented within a 4° aperture. The target stimulus remained on for 500 ms. When the stimulus disappeared, the fixation point also turned black, indicating the start of the response period. Subjects judged whether the target was more similar to naturalistic or spectrally matched noise by making a saccade to one of the choice targets. If the monkeys made a saccade to the correct choice target, they received a liquid reward. Responses to ambiguous stimuli were rewarded randomly.

We varied naturalness over a range from fully naturalistic (naturalness = 1) to fully phase-randomized (naturalness = 0). Stimuli were presented in random order. On each trial, naturalistic texture stimuli were drawn from 300 different possible samples within each naturalness condition. New samples were generated for each texture family once a week to prevent the animal from memorizing individual samples. However, on a subset of tasks in the ambiguous condition where naturalness = 0.5, we overrepresented a single sample to measure behavioral and neuronal covariability without the added variance from the presentation of different samples. Trials in which the subject did not maintain fixation within 0.6° of the fixation point were aborted. Data are reported from every session for which at least 100 trials were completed (230 in monkey 1, 222 in monkey 2).

We compared results from the texture discrimination task with previously published data where the same two monkeys performed an orientation discrimination task (Goris et al., 2017). Both monkeys were trained and then recorded from during the orientation discrimination task, and subsequently introduced to, trained on, and recorded from during the texture discrimination task. The logic and timing of the orientation discrimination task were identical to the texture discrimination task (Fig. 5c). In the orientation discrimination task, the choice targets were white lines (0.3° wide, 2.0° long) rotated –22.5° and 22.5° away from the discrimination boundary (which changed each session according to the preference of the recorded neuron). The target was a drifting grating and subjects judged the orientation of the stimulus relative to the discrimination boundary by making a saccade to one of the choice targets. See Goris et al. (2017) for further details.

### Analysis of behavioral response

We measured an observer’s sensitivity to the presence of naturalistic image structure by fitting the relationship between stimulus naturalness (ranging between 0 and 1) and probability of naturalistic choice with a psychometric function consisting of a lapse-rate and a cumulative Gaussian function, consistent with the methods of Goris et al. (2017). We defined the decision criterion and sensitivity as the mean and reciprocal of the standard deviation of the underlying Gaussian, respectively. We optimized the three parameters of the behavioral model (lapse rate, criterion, sensitivity) by maximizing their likelihood over each session’s data. Lapses were rare in most sessions, indicating both observers nearly always used the stimulus to inform their responses (Fig. 2c,d). To ensure this, we excluded a small number of sessions in which the lapse rate exceeded 15% from further analyses (7 out of 230 in monkey 1, 21 out of 222 in monkey 2).

### Analysis of neuronal response

We analyzed neural data within sessions where sufficient spike isolation was maintained throughout the entire session (153 out of 223 in monkey 1, 177 out of 201 in monkey 2). We summarized each neuron’s stimulus response with the number of spikes observed in a 500 ms window following response onset. For each cell, we determined the latency from stimulus presentation to response onset by maximizing the stimulus-associated response variance (Smith et al., 2005).

We estimated neuronal discrimination capabilities by fitting the relationship between stimulus naturalness and probability of “naturalistic” choice for an ideal observer with a cumulative Gaussian function. The ideal observer’s choices were obtained by applying a deterministic decision criterion to the responses of each neuron. To minimize bias in the ideal observer’s choices, the criterion was set to the median response to the ambiguous, 0.5 naturalness stimulus. We defined neuronal sensitivity as the reciprocal of the standard deviation of the cumulative Gaussian. We analyzed data from all neurons regardless of the level of sensitivity.

For each cell, we computed choice probability for the zero-signal stimulus and the neighboring stimulus conditions. As described previously (Britten et al., 1996), choice probability is calculated by performing an ROC analysis on the choice-conditioned neuronal responses to repeated presentations of a single stimulus. We classified a choice probability estimate as statistically significant if it fell outside of the central 95% of the expected null distribution, computed from 1000 randomly permuted data sets (Britten et al., 1996). The responses of many recorded neurons had very low sensitivity, rendering the sign of their choice probability (i.e. above or below 0.5) relatively uncertain. To remain consistent across different analyses, we determined the sign of choice probability estimates based on the the slope of a logistic regression analysis between the spike counts of individual neurons and an animal’s choices across different stimulus naturalness conditions (and excluding the ambiguous stimulus condition). The slope of this function reflects the strength and sign of each neuron’s tuning. To directly compare the relationship between spike counts and behavior in the presence and absence of variation in stimulus naturalness, we performed the same logistic regression analysis on spike counts only from the 0.5 naturalness condition. For comparisons across tasks, we performed the same analysis on previously collected data from the same two monkeys during an orientation discrimination task from Goris et al. (2017). In accordance with our approach to analyzing the texture discrimination data, we did not exclude any neurons from the dataset on the basis of weak orientation selectivity (monkey 1: 118; monkey 2: 116).

We computed average peristimulus time histograms by counting spikes in 1 ms windows for individual neurons, smoothing with a causal exponential filter with a time constant of 10ms, and averaging across all neurons. To compute the dynamics of choice probability we calculated the estimated choice probability for individual neurons within 100 ms time windows each shifted by 10 ms before averaging together these values across the population.

## Results

### Neuronal sensitivity to naturalistic visual structure in awake monkey V1 and V2

To study the neural basis of perception of naturalistic image structure, we created a set of model-based synthetic texture stimuli that have been well studied in perception and neurophysiological research (Portilla and Simoncelli, 2000; Freeman and Simoncelli, 2011; Freeman et al., 2013; Ziemba et al., 2016; Okazawa et al., 2015, 2017; Ziemba et al., 2018, 2019; Ziemba and Simoncelli, 2021). The model measures the marginal and joint statistics across the outputs of a simulated population of V1 simple and complex cells, in response to a photographic image of a natural texture (Fig. 1a; Portilla and Simoncelli (2000)). The same statistics are then measured from an input image of Gaussian white noise and iteratively adjusted until they match the statistics from the original image. If only the second-order, or spectral, statistics from the original image are imposed, the result is what we refer to as a “noise” image. If additional higher-order joint statistics across the output of V1-filters are matched to the original, the resulting “naturalistic” texture image often more closely resembles the original image (Fig. 1a; Portilla and Simoncelli (2000)). Starting the synthesis from a different input white noise image results in different samples of statistically matched texture. When the response over many samples is averaged, V1 neurons respond with similar firing rates to naturalistic and noise images, whereas V2 neurons generally respond with higher firing rates to naturalistic images (Freeman et al., 2013; Ziemba et al., 2018, 2019). These experimental observations were obtained from anesthetized macaque monkeys, and here we sought to test whether this distinction between V1 and V2 selectivity persists under conditions where subjects are awake and behaving.

**Figure 1.**
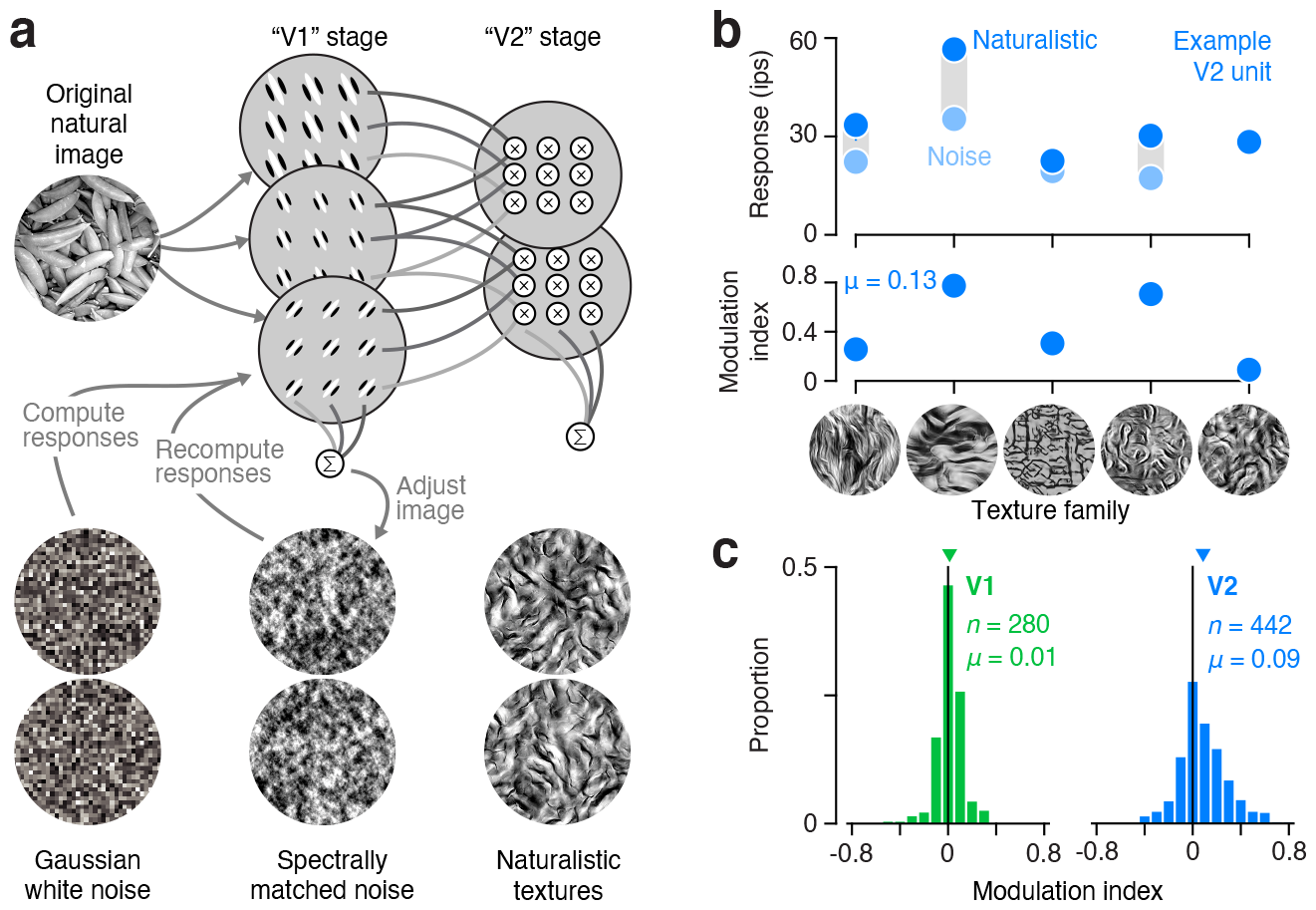
Tuning to “naturalness” in V1 and V2 of awake monkeys. (**a**) Schematic of texture synthesis procedure. An original black and white photograph is decomposed into the responses of a population of V1-like filters tuned to different orientations and spatial frequencies. Both the linear responses, and their squared energy, are computed, representing simple and complex cells, respectively. A second stage computes local correlations across the V1 responses tuned to different orientations, spatial frequencies, and positions and spatially averages them. To produce a synthetic texture, the model analyzes an image of Gaussian white noise and iteratively adjusts it to more exactly match the statistics of the original image. If the image is matched for the full set of correlations, we refer to it as a “naturalistic texture”, and if only matched for statistics capturing the spectral (second-order) content of the original image, we refer to it as “spectrally matched noise”. (**b**) Mean responses of an example V2 neuron to multiple samples of naturalistic (dark blue) and noise (light blue) images from five different texture families (top). Modulation index computed from these responses for five texture families (middle). (**c**) Distribution of modulation index values for a population of V1 (top) and V2 (bottom) neurons recorded from two awake, fixating macaque monkeys. Downward arrows indicate distribution mean, *μ*.

We installed a recording chamber with access to both V1 and V2 in two macaque monkeys trained to perform a texture discrimination task. During each recording session, we lowered an electrode into the visual cortex and isolated a single unit in either V1 or V2. We hand-mapped receptive fields to determine the receptive field center while the monkey fixated on a central spot. For subsequent measurements, the animal maintained fixation while several naturalistic and noise textures were shown, centered on the receptive field and presented within a four degree aperture. We presented multiple samples of naturalistic and spectrally matched noise images from five different texture families, each presented for 100 ms and separated by 100 ms of gray screen (Fig. 1b). We calculated a modulation index by dividing the difference in response to naturalistic and noise by the sum. This index captures the strength of preference for naturalistic images for each texture family (Fig. 1b, bottom). We selected the texture family that evoked the most discriminable responses between naturalistic and noise for the subsequent behavioral task, regardless of whether the neuron preferred naturalistic or noise stimuli. That is, we chose the texture family (out of five) most likely to reveal a relationship between the responses of the particular neuron under study and the subject’s behavior in the discrimination task.

Examining the average modulation index over all five texture families for the entire population of recorded neurons, we found a similar pattern to that observed in anesthetized macaque cortex (Fig. 1c). Specifically, there was little overall preference in the V1 population for naturalistic or spectrally matched noise stimuli and the average modulation index was near zero. In contrast, V2 neurons were mostly driven to higher firing rates by naturalistic stimuli, leading to more positive modulation indices. This pattern was consistent across both monkeys (monkey 1 V1: mean = 0.00, n = 137; monkey 1 V2: mean = 0.08, n = 250; monkey 2 V1: mean = 0.02, n = 143; monkey 2 V2: mean = 0.09, n = 192).

Unlike the average modulation index across textures in the original report recorded in anesthetized animals (Freeman et al., 2013), V1 neurons were significantly shifted toward positive values (P = 0.012; *t*-test on signed modulation; mean = 0.01, n = 280; Fig. 1c, left). However, here we used only the five texture families that yielded the highest average modulation index in V2 from the set of fifteen in the original study. Since the average modulation for each texture family was correlated across V1 and V2, the average modulation index in anesthetized V1 for these five textures was also significantly shifted toward positive values (P = 0.006; *t*-test on signed modulation; mean = 0.03, n = 102). There was no significant difference between average modulation index recorded previously from anesthetized V1 and here from awake V1 (P = 0.1; *t*-test on signed modulation). However, the modulation index in V2 was larger in the anesthetized monkey (mean = 0.22, n = 103) compared with awake V2 recordings made here (P < 0.001; *t*-test; mean = 0.09, n = 442; Fig. 1c, right). This may in part reflect regression to the mean, as the stimuli used here were chosen on the basis of the high V2 modulation index in the anesthetized experiments. Importantly, in both anesthetized and awake monkeys, V2 neurons had a significantly higher modulation index compared with those in V1 (P < 0.001; *t*-test).

### Naturalistic texture discrimination

In each recording session, after examining the tuning and determining the texture family yielding the highest sensitivity for the recorded neuron, monkeys performed a texture discrimination task, indicating the “naturalness” of a peripherally presented patch of texture (Fig. 2a). On each trial, after the animal attained stable fixation, we presented two choice targets to the left and right of fixation in the upper visual field. One choice target was a naturalistic texture matched to an original natural photograph for the joint statistics of the outputs of differently tuned V1-like filters (Portilla and Simoncelli, 2000). The other choice target was spectrally matched noise, lacking the higher-order statistics of the naturalistic texture. 500 ms after choice target onset, we presented a target stimulus centered on the receptive field of the simultaneously recorded single unit in V1 or V2. We generated the target stimulus by synthesizing intermediate textures whose statistics were linearly interpolated between the naturalistic and spectrally matched noise endpoints (Fig. 2b)(Freeman et al., 2013; Vacher et al., 2020). When no higher-order statistics were included in the synthesis (naturalness = 0) the stimulus was spectrally matched noise, and when the higher-order statistics were fully imposed (naturalness = 1) the stimulus resembled naturalistic texture. Stimuli with different levels of naturalness were synthesized using the same image seed, and this seed was randomized across trials so any particular spatial feature provided no cue for completing the task. Only discrimination of the higher-order statistics themselves allowed for high performance. The monkey indicated whether the target was more naturalistic or noise-like by making an eye movement to one of the choice targets following stimulus offset after 500 ms. Target stimuli above 0.5 naturalness were rewarded for a saccade to the naturalistic choice target, and those below 0.5 naturalness were rewarded for a saccade to the noise target.

**Figure 2.**
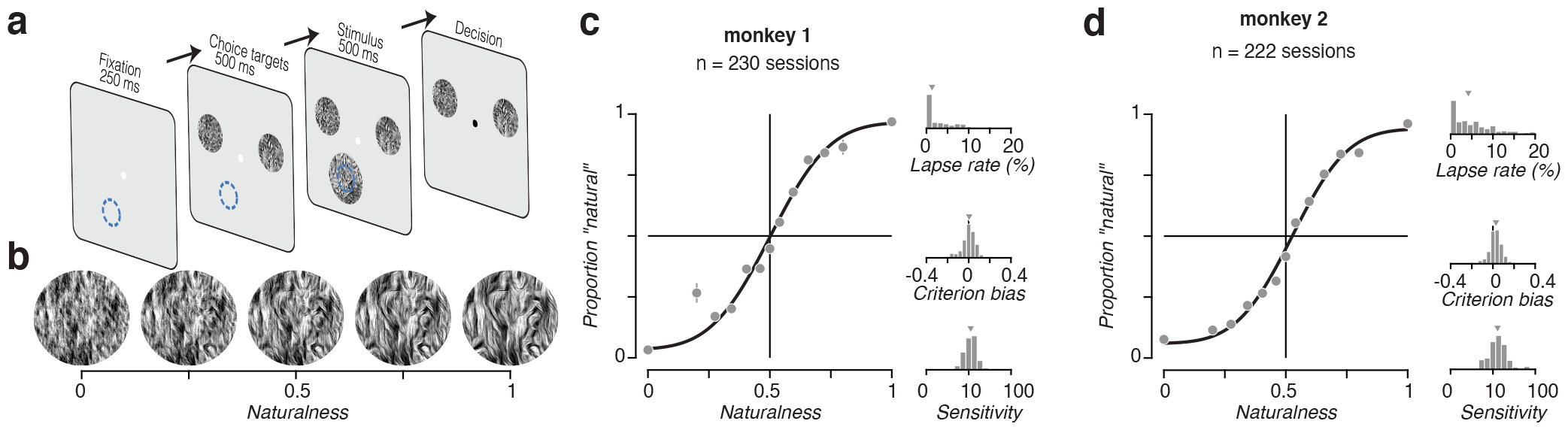
Naturalistic texture discrimination task. (**a**) After the subject acquired fixation for 250 ms, two choice targets appeared (one noise and one naturalistic). After another 500 ms, a stimulus was presented in the neuron’s receptive field (blue circle). Subjects judged whether the statistics of this stimulus were closer to noise or naturalistic texture. After a 500 ms presentation, the stimulus disappeared and the subject communicated their decision with a saccade toward one of the two choice targets. Rewards were given for correct answers, defined relative to naturalness value of 0.5, with stimuli at this boundary rewarded randomly. (**b**) Example stimuli along naturalness axis, for one texture family. (**c**) Behavioral performance of monkey 1 over many sessions. Left, average psychometric performance across all sessions. The points represent measured behavior. The line represents a fit of the signal detection theory model of behavior. Right top, distribution of guess rate over all sessions. Middle, distribution of bias parameter over all sessions. Bottom, distribution of sensitivity over all sessions. (**d**) Same as (c) but for monkey 2.

Both subjects performed the task well. When the target had a naturalness value of 0 or 1, performance was nearly perfect, and progressively declined as naturalness approached the experimenter-induced decision boundary at 0.5 (Fig. 2c,d). We fit the animals’ behavioral data for each session with a model in which choices arise from comparing a learned decision criterion to a noisy estimate of naturalness, with occasional “lapse” trials. Sensitivity, the slope of the resulting psychometric function at the fitted decision criterion, corresponds to the reciprocal of the fitted noise standard deviation. Both subjects had similar high sensitivity and little bias, despite having to learn the 0.5 naturalness boundary for each family (Fig. 2c,d).

### Comparing neuronal and perceptual sensitivity for naturalistic visual structure

Tuning for naturalness was, on average, modest in both V1 and V2. We assigned single units from each area according to their preference for naturalistic or spectrally matched noise images (see methods), and examined the average responses to all stimuli for each group. The groups were of similar size in V1, but in V2 there were more than twice the number of naturalistic preferring neurons (Fig. 3a). These naturalistic preferring V2 neurons on average fired *∼*15 impulses per second more for naturalistic stimuli compared with noise. This response differential was *∼*8 impulses per second for noise-preferring V2 neurons, and was *∼*6 in V1 neurons regardless of preference.

**Figure 3.**
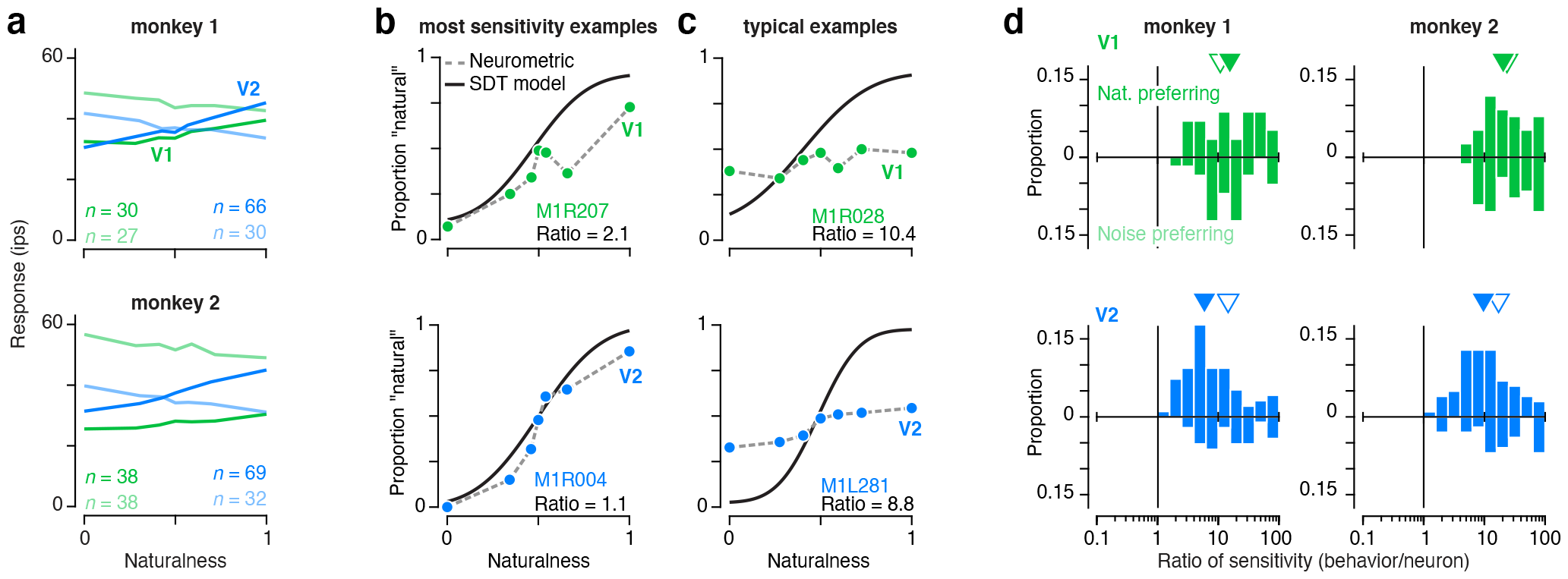
Comparison of neural and behavioral sensitivity. (**a**) Average naturalness tuning for populations of V1 (green) and V2 (blue) neurons preferring natural (dark) or noise (light) textures, plotted separately for monkey 1 (top) and monkey 2 (bottom). (**b**) Neurometric and psychometric functions obtained for sessions yielding the closest match between neuronal and behavioral sensitivity for V1 (top) and V2 (bottom). Black line represents the signal detection theory model fit to behavioral responses. Colored symbols represent ideal observer analysis applied to the responses of single neurons. Neuronal sensitivity was obtained by fitting the signal detection theory model to these neurometric data. (**c**) Neurometric and psychometric functions recorded from sessions yielding ratios of neuronal to behavioral sensitivity that were more typical of the V1 (top) and V2( bottom) populations. (**d**) Distribution of the ratio between single neuron and behavioral sensitivity. Upright distributions represent naturalistic-preferring neurons, and upside down distributions represent noise-preferring neurons. Ordinate axes in each panel indicate proportion across the entire population, and downward arrows indicate the median ratio for naturalistic- (filled) and noise-preferring (empty) neurons. In both monkeys, V2 neurons (bottom) were significantly closer to the sensitivity of behavior than V1 neurons (top), and naturalistic-preferring neurons in V2 were significantly closer to the sensitivity of behavior than noise-preferring neurons. There was no different between naturalistic- and noise-preferring neurons in V1.

Although both subjects learned to perform the task well and showed consistent sensitivity across sessions, the sensitivity of single neurons to naturalness varied widely. To estimate neuronal sensitivity, we performed an ideal observer analysis on the distribution of spike counts to different levels of naturalness. We applied a decision criterion at the median spike count elicited by stimuli at 0.5 naturalness, and took any response above this criterion as a neuronal “decision” to report natural. When plotted as a function of stimulus naturalness, the proportion of natural responses traces out a neurometric function (Fig. 3b,c). The slope of this function serves as a measure of the sensitivity of the neuron to changes in naturalness, and can be directly compared to sensitivity measured from the psychometric function. Examining the most sensitive neurons demonstrates that on occasion neuronal sensitivity approached that of the animal’s choices, especially in V2 (Fig. 3b). However, more typically, the sensitivity of the monkey far exceeded that of single neurons in both V1 and V2 (Fig. 3c).

Both subjects showed a similar pattern in the relationship between neuronal and behavioral sensitivity. Sensitivity in V2 was significantly closer to behavior than in V1 (Fig. 3d; P < 0.001, Wilcoxon rank sum test). On average, V1 neurons were 18 times less sensitive than the animal’s behavior while V2 neurons were about 10 times less sensitive than behavior. In V1, there was no difference between neurons that preferred naturalistic to those that preferred noise (median = 17.0 versus 20.4; P = 0.93, Wilcoxon rank sum test). However, in V2 in both monkeys, neurons that preferred naturalistic images had sensitivity significantly closer to behavior (median = 7.4 versus 17.1; P < 0.001, Wilcoxon rank sum test). Although we have previously hypothesized that V2 neurons may play a role in the perception of naturalistic texture (Freeman et al., 2013), this seven-fold difference between behavioral and neuronal sensitivity is substantially larger than in many studies that have compared behavior with a potential neural correlate (Britten et al., 1992; Nienborg and Cumming, 2006, 2014; Goris et al., 2017). However, unlike most of these studies, we had limited ability to optimize the texture discrimination task to the particular selectivity of the recorded neuron because of the complexity of the space of possible naturalistic textures. A previous study that recorded populations of V1 and V4 units while monkeys performed a fixed orientation discrimination task found average behavioral to neuronal sensitivity ratios greater than 20 (Jasper et al., 2019). In contrast, when a monkey performs an orientation task adjusted to the preference of a recorded V1 neuron, average behavioral to neuron sensitivity ratios are close to one (Nienborg and Cumming, 2014; Goris et al., 2017). This comparison suggests that if we performed a more comprehensive characterization of the naturalistic texture selectivity of V2 neurons and optimized the task stimuli accordingly, neuronal sensitivity might be much closer to behavior.

### Inconsistent choice-correlated activity across subjects and tasks in V1 and V2

We wondered whether we could find evidence for the potential participation of our recorded population in the formation of perceptual decisions about naturalistic texture despite the discrepancy between neuronal and perceptual sensitivity. In particular, we wondered whether the increased sensitivity of V2 neurons would manifest in a greater tendency to predict perceptual decisions on a trial-by-trial basis. To examine this, we computed the “choice probability” for the responses of each neuron to the ambiguous, 0.5 naturalness condition (Fig. 4a) (Britten et al., 1996). This quantity corresponds to the probability that a neuron fires more spikes preceding a behavioral decision associated with its preferred stimulus. A choice probability of 0.5 indicates that a neuron’s response contains no information about choice and a value of 1 indicates perfect prediction of choice.

**Figure 4.**
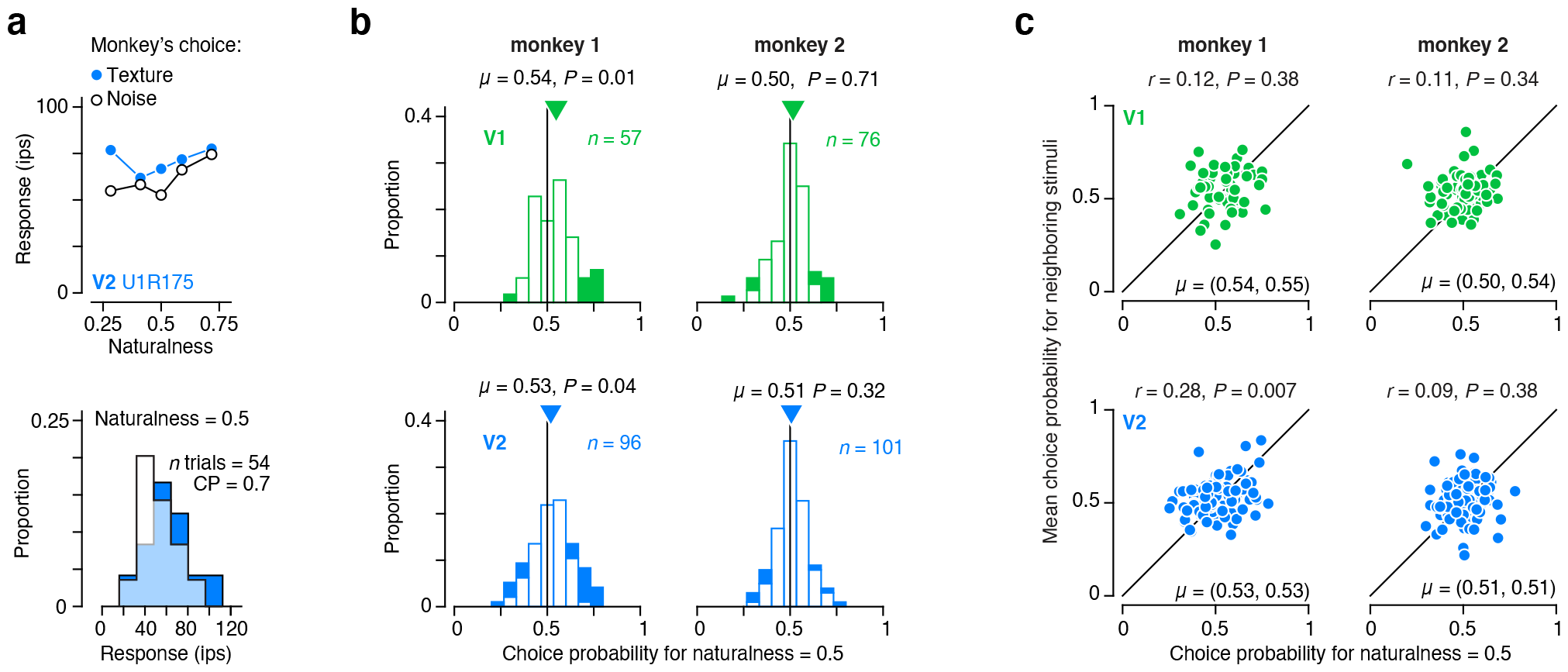
Choice probability for naturalistic texture discrimination. (**a**) Mean responses from an example V2 neuron conditioned on whether the monkey chose “texture” or “noise” on a given trial (top). Distribution of responses when naturalness = 0.5 conditioned on the behavioral choice (bottom). (**b**) Distribution of choice probability for naturalness = 0.5 for V1 (top) and V2 (bottom) neurons recorded from monkey 1 (left) and Monkey 2 (right). Filled bars represent neurons with choice probability significantly different from 0.5 as determined by permutation test. Mean choice probability in V1 and V2 was significantly larger than 0.5 in monkey 1 (P < 0.05; *t*-test), but not monkey 2 (P > 0.05; *t*-test). (**c**) Mean choice probability for the two stimulus conditions flanking naturalness = 0.5, plotted against choice probability for naturalness = 0.5 for V1 (top) and V2 (bottom) neurons recorded from monkey 1 (left) and monkey 2 (right).

The two monkeys differed in the pattern of their choice-correlated activity. In monkey 1, there were many neurons with strong choice-correlated activity (Fig. 4a) and the mean choice-probability in both V1 and V2 was significantly larger than 0.5 (Fig. 4b, left). However, we found no evidence that mean choice probability differed from 0.5 in monkey 2 in either V1 or V2 (Fig. 4b, right). While the inconsistency between the two monkeys is not unprecedented, we were surprised to find little difference in the strength of choice-correlated activity in V1 and V2, given the very weak neuronal sensitivity to texture naturalness in V1 neurons compared with V2. To examine the consistency of this choice-correlated activity, we also calculated the average choice probability for the two stimulus conditions adjacent to the ambiguous, 0.5 naturalness condition. For monkey 1, we found no significant correlation in choice probability across conditions in V1 (*r* = 0.12; P = 0.38), but we found a significant relationship in V2 (*r* = 0.28; P = 0.007; Fig. 4c). Despite this, choice probability was still significantly larger than 0.5 in V1 for neighboring conditions (mean = 0.055; P = 0.001; *t*-test). For monkey 2, we found no significant correlation in either V1 (*r* = 0.11; P = 0.34) or V2 (*r* = 0.09; P = 0.38). However, choice probability was actually significantly larger than 0.5 exclusively when computed from the adjacent conditions of V1 in monkey 2 (mean = 0.54, P < 0.001; *t*-test). In summary, choice-correlated activity was a stable and consistent feature of neural responses only in V2 of monkey 1.

The stimuli in our texture discrimination task differ from most previous experiments for which significant choice-correlated activity has been observed. While most studies use dynamic, noisy stimuli, we presented a single, static sample of texture for 500 ms during each trial (but see (Kosai et al., 2014)). Our previous results indicate that differences in statistically matched texture samples account for a substantial amount of neuronal variability in V1 and V2 (Ziemba et al., 2016)). To examine whether texture sample variability might drive our choice probability observations, we presented different texture samples multiple times over the course of a session. We calculated a grand choice probability value by combining neural responses across multiple presentations of the same texture sample that led to different behavioral responses. The results of this analysis were similar to our previous observations for both monkey 1 (V1: Mean = 0.54, P = 0.003, *t*-test; V2: Mean = 0.53, P = 0.04), and monkey 2 (V1: Mean = 0.50, P = 0.81, *t*-test; V2: Mean = 0.52, P = 0.04).

Given the lack of a coarse relationship between neuronal sensitivity and the presence of choice-correlated activity across areas, we developed a more nuanced way of examining this relationship across all neurons (Zaidel et al., 2017). Combining the logic of choice probability with the power of logistic regression instead of ROC analysis, we applied the same method to quantify the strength of both sensitivity and choice-correlated activity. For each neuron we used logistic regression to predict the choice of the animal based on the observed spike count on each trial. First, we found the slope coefficient when estimating the model across stimulus conditions (but excluding the ambiguous 0.5 naturalness condition; Fig. 5a, left). Given the lawful behavioral performance, this across-condition slope essentially measures how strongly changes in stimulus naturalness affect the response of a neuron and the behavioral response of the animal. The sign of the slope indicates whether higher naturalness tends to increase or decrease responses, and the magnitude indicates the strength of the relationship between spikes and behavior. Second, we independently fitted the slope coefficient when estimating the model using only responses elicited by the ambiguous 0.5 naturalness stimulus condition (Fig. 5a, right). This within-condition slope estimates how changes in the response of the neuron predict behavioral responses without variation in stimulus naturalness — analogous to choice probability. Like the across-condition slope, the sign of the slope indicates whether a behavioral choice for naturalness tends to increase or decrease responses, and the magnitude indicates the strength of this relationship.

**Figure 5.**
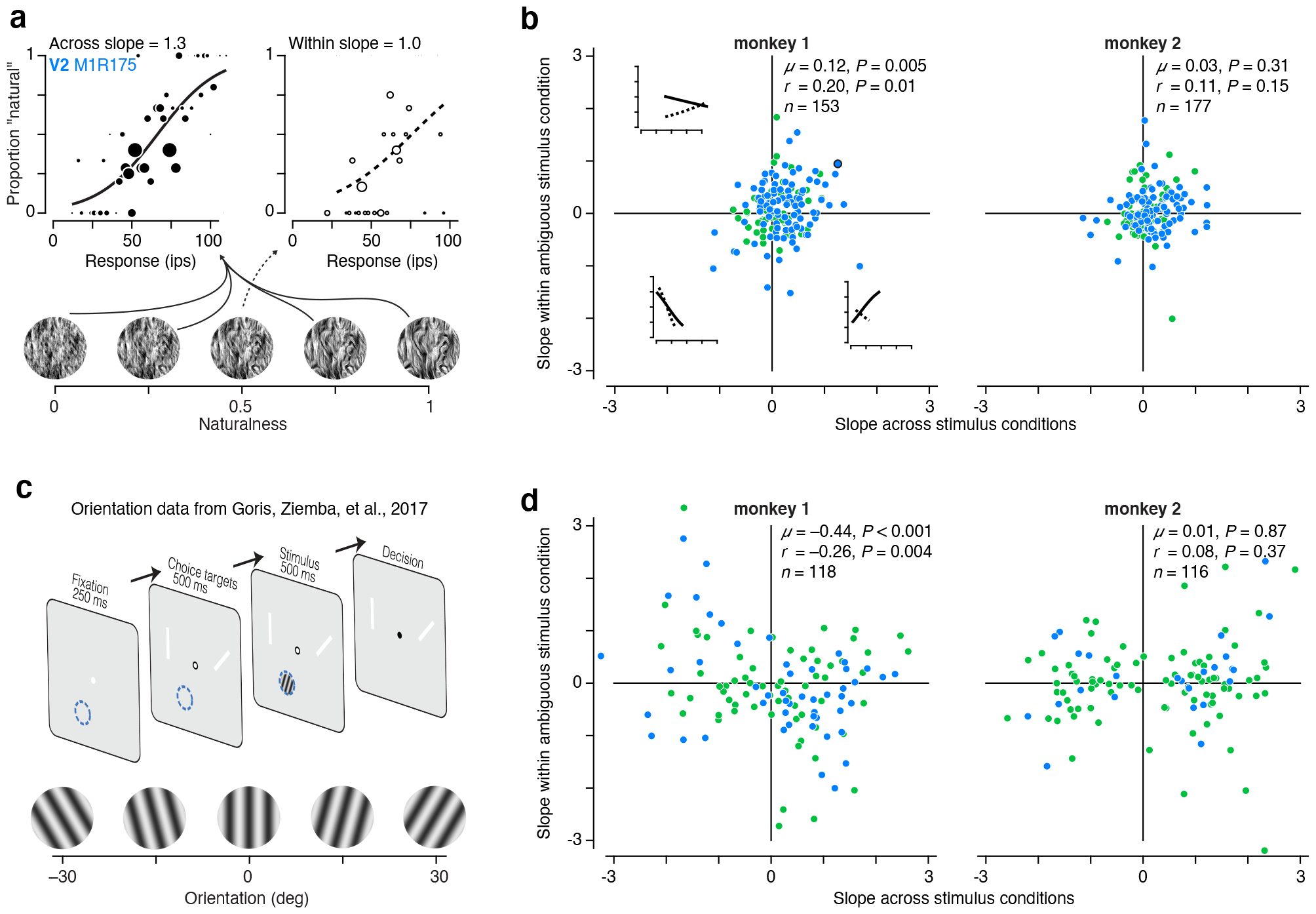
Trial-by-trial relationship between behavioral choice and neural activity. (**a**) The choices of the monkey plotted against responses of an example V2 neuron for all conditions where naturalness ≠ 0.5 (left), and when naturalness = 0.5 (right). The diameter of each symbol indicates the number of responses of that magnitude observed. Lines represent the fit of a logistic regression analysis performed separately across (left, solid) and within (right, dashed) stimulus conditions. (**b**) The slope of the logistic regression analysis performed across conditions plotted against the analysis performed within conditions for monkey 1 (left) and monkey 2 (right). Insets show example across (solid) and within (dashed) fits within quadrants 2-4. Symbol outlined in black indicates example neuron shown in (a). (**c**). Previously reported orientation discrimination task. (**d**) Slope of logistic regression across vs within conditions (as in (b)) for the orientation discrimination task (same two monkeys).

If these neurons informed perceptual choices or were involved in the perceptual inference process, we would expect the sign of the relationship between the neural and behavioral response to remain the same in both the presence and absence of stimulus variation (as with the example neuron in Fig. 5a). This would mean that if we plot the across-condition slope against the within-condition slope we should expect most neurons to fall in quadrants one and three. We tested for this expected pattern of choice-correlated activity by examining the within-condition slope multiplied by the sign of the across-condition slope. This quantity will be above zero if the within- and across-condition slopes generally match in their sign. We found evidence for this expected pattern in monkey 1 (Fig. 5b, left; V1/V2 mean = 0.12, P = 0.005, *t*-test; V1 mean = 0.15, P = 0.02; V2 mean = 0.10, P = 0.08), but not monkey 2 (Fig. 5b, right; V1/V2 mean = 0.03, P = 0.31, *t*-test; V1 mean = 0.02, P = 0.77; V2 mean = 0.04, P = 0.26). However the difference between the two monkeys did not quite reach conventional statistical significance (P = 0.09, *t*-test). We also found a significant correlation between within- and across-condition slopes in monkey 1 (V1/V2 r = 0.20, P = 0.01; V1 r = 0.25, P = 0.07; V2 r = 0.18, P = 0.07), but not monkey 2 (V1/V2 r = 0.11, P = 0.15; V1 r = 0.11, P = 0.33; V2 r = 0.10, P = 0.34).

Given these apparent differences in the pattern of choice-correlated activity across monkeys, we wondered whether these patterns might at least be a stable feature across time and results from different tasks in these animals. Both monkeys previously participated in experiments where they discriminated the orientation of a drifting grating while we recorded single unit activity in V1 and V2 (Fig. 5c) (Goris et al., 2017). The orientation discrimination task followed an identical logic and temporal structure. Choice targets were white lines oriented 45° apart. The orientation midway between the two choice targets communicated the discrimination bound, and we rewarded monkeys for making a saccade to the target whose orientation most closely matched the stimulus orientation. This task structure allowed us to move the discrimination bound to an orientation that matched the steepest part of a recorded neuron’s orientation tuning curve in addition to optimizing the stimulus diameter, spatial frequency, and drift rate to elicit the largest response. As in the texture discrimination task, we presented the ambiguous orientation at the discrimination bound twice as often as other stimuli. This allowed us to perform the same logistic regression analysis on these data — computing the across-condition and within-condition slope and comparing them. The results examined in this way match our original observations using choice probability (Goris et al., 2017), but are strikingly different than the results from the texture discrimination task in monkey 1 (Fig. 5d, left).

First, across-condition slopes were much higher for orientation than for texture, which is to be expected given our ability to optimize the task for the neuron’s tuning in the orientation discrimination task. More importantly, the within-condition slope multiplied by the sign of the across-condition slope was significantly negative (Fig. 5d, left; V1/V2 mean = -0.44, P < 0.001, *t*-test; V1 mean = -0.44, P = 0.008; V2 mean = -0.43, P < 0.001), and the correlation between within and across condition slopes was negative (r = -0.26, P = 0.004; V1 r = -0.22, P = 0.08; V2 r = -0.36, P = 0.009). The within-condition slope multiplied by the sign of the across-condition slope was significantly different for monkey 1 during the orientation discrimination task compared with the texture discrimination task (P < 0.001, *t*-test). However, as in the texture discrimination task, there was no evidence for choice-correlated activity in monkey 2 in either the average within-condition slopes (Fig. 5d, right; V1/V2 mean = 0.01, P = 0.87, *t*-test; V1 mean = -0.03, P = 0.74; V2 mean = 0.15, P = 0.33) or the correlation between across- and within-condition slopes (r = 0.08, P = 0.37; V1 r = 0.01, P = 0.94; V2 r = 0.34, P = 0.09).

We were struck by this complex pattern of results. In the texture task, despite much higher neuronal sensitivity in V2, there was no tendency for choice-correlated activity to be stronger than in V1. Further, in the orientation task, neuronal sensitivity was much higher than in the texture task, but choice-correlated was either similarly low or systematically misaligned with stimulus preference compared with activity in the texture task. This pattern of inconsistency across observers within the same task and across tasks in the same observer suggests that the presence of choice-correlated activity is not a reliable indicator of the participation of a neural population in the perceptual inference process (Goris et al., 2017; Jasper et al., 2019; Zaidel et al., 2017; Yu and Gu, 2018; Krishna et al., 2021; Zhao et al., 2020; Quinn et al., 2021; Lange et al., 2023; Levi et al., 2023). Instead, choice-correlated activity may likely arise from feedback to sensory areas reflecting a mixture of factors not necessarily related to decision formation (Goris et al., 2017; Quinn et al., 2021; Laamerad et al., 2023)

### Dynamics of neuronal responses in V1 and V2

We wondered whether an analysis of the temporal evolution of responses might reveal the origin of texture selective and choice-correlated activity in V1 and V2. We first examined the temporal form of the average firing rate across our population of V1 and V2 neurons (Fig. 6a). In both areas, there was a prominent initial transient in firing rate following stimulus onset. The transient amplitude was preserved across different naturalness conditions, indicating that this initial portion of the response contained little stimulus selectivity. To quantify the dynamics of population selectivity, we computed the variance of the average firing rate across conditions, yielding a measure of the stimulus induced variance.

**Figure 6.**
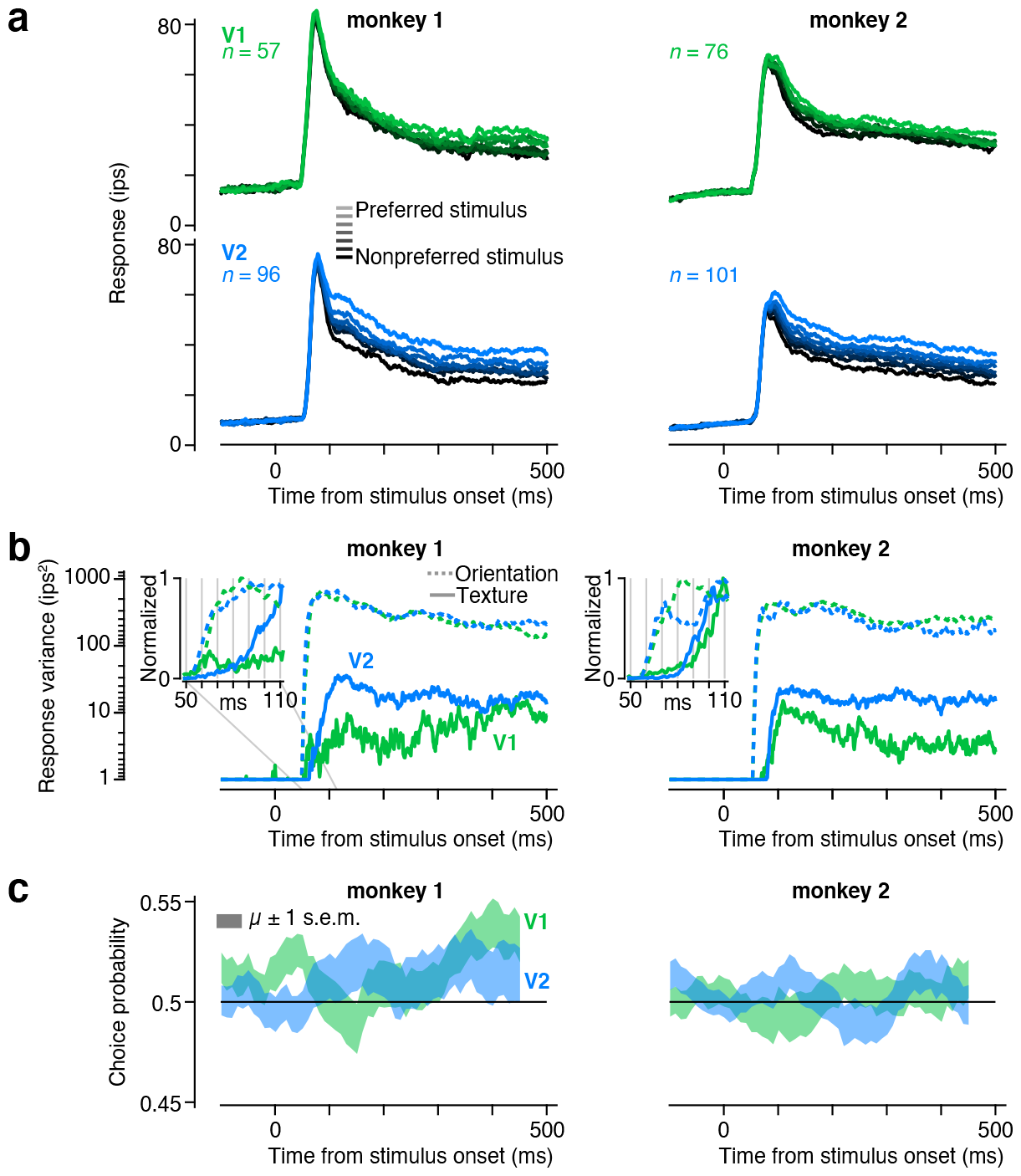
Dynamics of neuronal sensitivity and choice-correlated activity. (**a**) Aver-age response dynamics in V1 (top) and V2 (bottom) for monkey 1 (left) and monkey 2 (right). The stimulus ordering was reversed for noise-preferring neurons before averaging, so that the lightest line in each panel represents responses to the preferred stimulus. (**b**) The variance of the average response over stimulus conditions in the texture discrimination task (solid lines) and the orientation discrimination task (dashed lines). Traces are plotted with a minimum value of 1 *ips*^2^. Insets expand the time window 50-110 ms after response onset and show all response variance values divided by the maximum value reached within the 500 ms stimulus window. (**c**) Dynamics of choice probability. We computed choice probability in 100 ms windows and plot the mean *±* standard error in the shaded region.

The dynamics of this selectivity measure differed markedly in V1 and V2 (Fig. 6b). For monkey 1, The stimulus-induced variance was weak in V1 and developed only gradually, peaking over 400 ms after stimulus onset (Fig. 6b, left). For monkey 2, this measure in V1 peaked earlier but then decayed (Fig. 6b, right). In contrast, for both monkeys, variance in V2 rose rapidly (within 100 msec) to a higher level than in V1, and this was largely maintained throughout the stimulus period. These differences between V1 and V2 were specific to naturalistic texture stimuli. When we examined the time course of orientation-induced variance, there was no prominent difference in either overall selectivity or the dynamics of selectivity between V1 and V2 (Fig. 6b, dashed lines). However, even in V2, the orientation-induced variance was more than an order of magnitude larger than that induced by varying naturalness.

Interestingly, although selectivity for naturalness peaked earlier in V2 than in V1, the peak was quite delayed with respect to response onset (Fig. 6b). This reflected the presence of a mostly non-selective transient response in the population firing rate (Fig. 6a). This pattern differed from that seen for orientation, which reached a peak value 20-30 ms earlier than for naturalness in both areas (Fig. 6b, insets). Thus, while orientation information appears immediately in the firing rate of individual neurons, the first spikes in V2 don’t appear to carry information about the naturalness of the stimulus.

Given the differences in the dynamics of selectivity between V1 and V2, we wondered whether the dynamics of choice-related activity would also differ between the two areas. We computed choice probability within 100 ms windows shifted by 10 ms for each individual neuron and then averaged the result. In monkey 2, choice probability was maintained around 0.5 throughout the trial in both V1 and V2. However, we found that choice-related activity was generally weak but evolved over a somewhat different time course in V1 and V2 of monkey 1 (Fig. 6c, left). In V1, choice probability was near chance for the first 300 ms following response onset, then began to rise and peaked just before stimulus offset at 500 ms. This temporal profile resembled the lack of early selectivity in V1, as well as the delayed onset of choice-correlated activity in both V1 and V2 observed in monkey 1 performing the orientation discrimination task (Goris et al., 2017). However, in V2, choice probability grew to be larger than 0.5 soon after response onset, peaking at about 150 ms post-stimulus and remaining roughly constant thereafter. This matches the earlier onset of choice-correlated activity observed in V2 compared with V1 for orientation discrimination (Goris et al., 2017), but without the delayed onset expected for a feedback signal.

### Dissociation of anticipatory signals and choice-correlated activity across subjects and tasks

The sequence and timing of events in both the orientation and texture discrimination tasks were identical across all trials, meaning that stimulus onset was always predictable (Fig. 2a, Fig. 5c). Some neurons anticipated the appearance of the stimulus in their receptive field through a gradual increase in their firing rate during the fixation period leading up to stimulus onset (Fig. 7a). Given the lack of external events during this period, this anticipatory signal must be internally generated. When these monkeys performed the orientation discrimination task, these anticipatory signals seemed to be related to choice-correlated activity (Goris et al., 2017). Both anticipatory and choice-related signals were present in neural activity recorded from monkey 1, but absent in monkey 2, suggesting a possible shared origin in feedback to visual cortex (Goris et al., 2017). However, we found that anticipatory signals were similarly unstable but unrelated to the changes in choice-correlated activity we observed across tasks.

**Figure 7.**
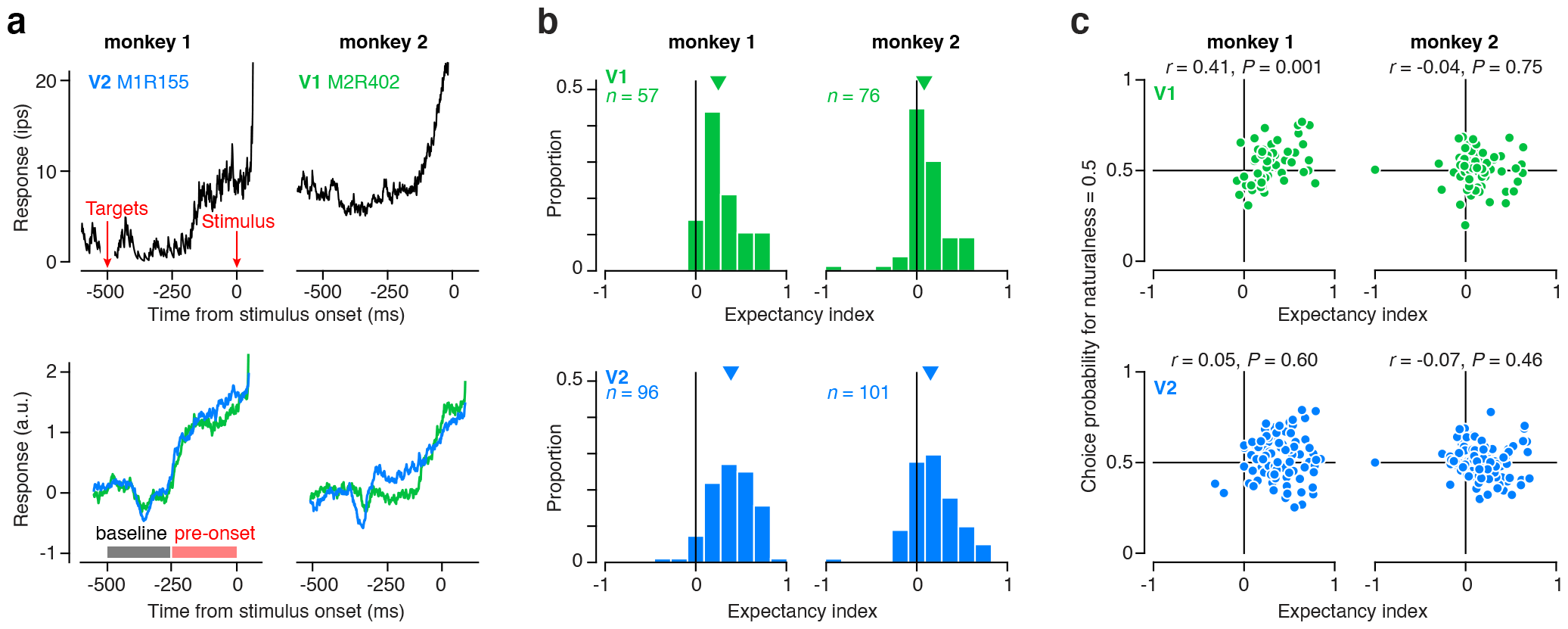
Analysis of anticipatory signals in V1 and V2. (**a**) Top, Response dynamics in anticipation of stimulus onset for two example neurons, recorded from monkey 1 (left, V2) and monkey 2 (right, V1). Peristimulus time histograms were created by averaging spike trains convolved with an exponential filter (tau = 10 ms). Bottom, Mean response dynamics in anticipation of stimulus onset for a population of V1 (green) and V2 (blue) neurons, recorded from monkey 1 (left) and monkey 2 (right). Before averaging, responses of each neuron were z-scored by their response in the baseline period. Filled bars represent the baseline and pre-onset intervals used to compute the expectancy index. (**b**) Distribution of the expectancy index for a population of V1 (green) and V2 neurons (blue), recorded from monkey 1 (left) and monkey 2 (right). (**c**) Choice probability for naturalness = 0.5 plotted against expectancy index for a population of V1 (top) and V2 (bottom) neurons, recorded from monkey 1(left) and monkey 2 (right).

There was significant anticipatory activity in the responses of neurons recorded from monkey 2 during the texture discrimination task, but no such activity during orientation discrimination (Fig. 7a, bottom; Goris et al. (2017)). In contrast, anticipatory signals were similar between the two tasks in monkey 1 (Goris et al., 2017). Activity in both areas in both monkeys dropped about 100 ms after the onset of the choice targets, likely due to suppressive effects of stimuli far outside the receptive field (Bair et al., 2003). Soon after returning to baseline, neural activity from monkey 1 began to rapidly rise in anticipation of the stimulus, whereas the onset of this anticipatory activity was about 100 ms delayed in monkey 2 (Fig. 7a, bottom). This difference somewhat mirrors the general delay in responses to texture stimuli in monkey 2 relative to monkey 1 (Fig. 6).

To quantify anticipatory signals, we computed an expectancy index as the difference in response between the first (baseline) and second half (pre-onset) of the fixation period and divided by the sum (Fig. 7a). This index ranges between -1 and 1 with positive values indicating an increased response in anticipation of the stimulus. For both animals and areas, the expectancy index was significantly positive (Fig. 7b; monkey 1 V1 median = 0.25, p < 0.001; monkey 1 V2 median = 0.39, p < 0.001; monkey 2 V1 median = 0.08, p < 0.001; monkey 2 V2 median = 0.15, p < 0.001, Wilcoxon signed rank test). The expectancy index was significantly larger in V2 compared with V1 in monkey 1 (p = 0.002, Wilcoxon rank sum test), and marginally larger in monkey 2 (p = 0.08, Wilcoxon rank sum test). Most interestingly, the expectancy index computed here for monkey 2 during the texture discrimination task was significantly larger than that found for the orientation discrimination task in both V1 (p < 0.001) and V2 (p = 0.004, Wilcoxon rank sum test; Goris et al. (2017)).

Based on the previous orientation discrimination experiments we suggested a link between choice-related activity and expectancy signals in visual cortex (Goris et al., 2017). However, when we collected neural data from these same animals retrained to perform a texture discrimination task, this link was broken. The sign of choice-related activity flipped in monkey 1, but the presence and strength of anticipatory signals remained largely unchanged. In contrast, data from monkey 2 showed no choice-related activity in either task, but significant anticipatory activity developed only in the texture discrimination task. At the single neuron level, there was a trend for neurons with stronger expectancy index to have higher choice probability in monkey 1 (Fig. 7c; across all neurons: r = 0.15, p = 0.06; V1: r = 0.41, p = 0.001; V2: r = 0.05, p = 0.6), but no evidence for such a trend in monkey 2 (across all neurons: r = –0.05, p = 0.47; V1: r = –0.04, p = 0.75; V2: r = –0.07, p = 0.46). This pattern is similar to that observed during the orientation experiments but with an opposite sign and a stronger relationship in V1 compared with V2 in monkey 1 (Goris et al., 2017). Thus, nonsensory, modulatory inputs are likely to be the common origin of both choice-related and anticipatory signals in visual cortex, but their presence and relationship can be decoupled and inconsistent across different tasks and observers.

## Discussion

Our experiments reveal functional differences between V1 and V2 neurons during the performance of a naturalistic texture discrimination task, but also highlight the limitations of neural signatures previously considered to reflect the participation of a neural population in perceptual decisions. We found greater sensitivity in V2 neurons for the higher order structure present in natural texture images, replicating our previous findings recorded under anesthesia (Freeman et al., 2013; Ziemba et al., 2016, 2019). We further show that despite the increased sensitivity from V1 to V2, V2 neurons are still less sensitive than behavior on average. This finding indicates that although sensitivity to naturalistic image structure first emerges in V2, it is likely further consolidated downstream (Arcizet et al., 2008; Rust and Dicarlo, 2010; Okazawa et al., 2015, 2017). V4 neurons are selective for the higher-order statistics present in our naturalistic stimuli (Okazawa et al., 2015), and a direct comparison suggests this selectivity is greater than for V2 neurons (Okazawa et al., 2017). Human neuroimaging results also suggest that areas downstream of V2, contain similar or increased sensitivity to complex image structure (Movshon and Simoncelli, 2015; Kohler et al., 2016; Henderson et al., 2023).

We observed a markedly different temporal emergence of sensitivity to naturalistic statistics compared with our previous work. This may reflect the effect of anesthesia on subcortical and cortical responses (Alitto et al., 2011; Jazayeri et al., 2012). In anesthetized recordings there was a gradual increase in responses to a stimulus and no initial transient (Freeman et al., 2013), potentially reflecting slower dynamics within individual neurons or more variability in the response latency across neurons (Jazayeri et al., 2012). Sensitivity to naturalistic statistics was also present at response onset and gradually developed over the course of tens of milliseconds under anesthesia (Freeman et al., 2013). Here, there was a significant delay in the onset of naturalistic sensitivity. This observation, along with previous findings that the onset of surround-enhanced sensitivity is delayed (Ziemba et al., 2018), suggests that intracortical processing within V2 (or potentially feedback from higher areas) plays a significant role in establishing selectivity for naturalistic image structure. The dynamics we report here also differ from the previously reported fast onset of “shape selectivity” recorded in awake V2 (Hegdé and Van Essen, 2004). However, this form of selectivity does not diverge markedly from that in V1, and so may be inherited (Hegdé and Van Essen, 2007). Along these lines, the dynamics of naturalistic sensitivity differ strikingly from the emergence of orientation selectivity (Goris et al., 2017), a property known to be influenced largely by linear filtering of feedforward inputs (Priebe and Ferster, 2012; Goris et al., 2015). In contrast, a functional account of naturalistic sensitivity requires two stages of processing in V2 (analogous to models of complex cells in V1), and the slow emergence of selectivity may reflect the dynamics of this computation (Freeman et al., 2013).

Although we found higher sensitivity for V2 neurons compared with V1, average sensitivity in V2 neurons was still far from behavioral performance. Many studies have found a tight correspondence between neuronal and behavioral thresholds (Britten et al., 1992; Prince et al., 2000; Uka and DeAngelis, 2006; Nienborg and Cumming, 2006, 2014), more clearly indicating a role for the responses of a particular brain area in supporting a particular perceptual behavior. However, each of these studies, as well as our own orientation discrimination experiment, involved tailoring the target stimulus and discrimination bound to the tuning preference of each individual neuron. This procedure gives the single neuron the best chance to match behavioral performance. We currently do not have a detailed understanding of the space of naturalistic statistics and therefore cannot perform comprehensive tuning experiments along meaningful axes of selectivity (although see Okazawa et al. (2015) for an example of adaptively searching the parameter space). Instead, we are limited to characterizing the differential response for a predefined set of texture families (for which the animal has been previously trained) and performing subsequent behavioral experiments using the most discriminable texture family for the recorded neuron.

The naturalistic sensitivity of V2 neurons is comparable to previous studies that have not optimized stimuli to the tuning preferences of recorded neurons (Purushothaman and Bradley, 2005; Gu et al., 2008, 2007; Shiozaki et al., 2012; Jasper et al., 2019). Average neuronal sensitivity in these studies can range from 3 to 20 times less sensitive than behavior. Most studies observe a pattern of wide distribution over neuronal sensitivity, with the most sensitive neurons approaching or overtaking behavioral sensitivity. This is consistent with our results in V2. However, V1 neuronal sensitivity was very weak and the most sensitive neuron was still two times less sensitive than behavior.

Many studies, both those that optimize stimuli for single neurons and those that don’t, have found significant choice-related activity. However, the interpretation of this observation has grown murkier over the last decade. It was initially suggested that response noise in sensory neurons causally affected the downstream decision process, manifesting as correlations between single sensory neurons and behavior (Britten et al., 1996; Shadlen et al., 1996; Parker and Newsome, 1998; Pitkow et al., 2015). Recent results have instead posited a feedback origin of such correlations, whereby the outcome of a perceptual decision is relayed to early sensory neurons (Nienborg and Cumming, 2009; Nienborg et al., 2012). The temporal emergence of choice probability has been important in disentangling contributions from feedforward versus feedback sources. Here we report two different temporal profiles of choice-related activity in V1 and V2 of one monkey. The dynamics of choice-related signals in V2 are similar to many previous reports in the literature, which show an early onset of significant choice probability that remains roughly constant throughout the stimulus period (Britten et al., 1996). Even recent theoretical work that posits a strong feedback component for choice probabilities finds that this flat temporal choice probability profile reflects an early feedforward contribution to perceptual decisions (Haefner et al., 2014; Wimmer et al., 2015; Haefner et al., 2016). Dynamics in V1 are less consistent with previous observations, and strongly suggest a feedback origin (Nienborg and Cumming, 2009).

As signals propagate along the cortical hierarchy, they encode increasingly complex aspects of the sensory environment and likely have a more direct relationship with perceptual experience. Over the past seven decades, visual neuroscience has been very successful in developing experimental methods that illuminate the first part of this statement. This has taken the form of characterizing the emergence of novel stimulus selectivities along the hierarchy (as replicated here) and of stimulus-response models that predict neural responses to arbitrary stimuli. However, we (as a field) have struggled to develop rigorous and robust experimental procedures that uncover the relationship between neuronal activity and perception. There have been some notable successes. For example, direct electrical micro-stimulation of sensory cortex can bias perceptual reports, suggesting that the stimulated neurons inform perceptual experience (Salzman et al., 1990; Yu and Gu, 2018). But this causal approach is limited in scope as it requires a coarse-scale organization of stimulus preference to perturb neural population activity on the “natural” manifold (Jazayeri and Afraz, 2017). Correlational approaches that characterize the relationship between neuronal and perceptual variability are more broadly applicable, but these studies have yielded inconsistent results. In some studies, the structure of choice-correlated activity is in line with simple feedforward hypotheses about the contribution of neural activity to perceptual decisions, meaning that suitably tuned neurons exhibit the strongest association with behavior (Britten et al., 1996; Dodd et al., 2001; Nienborg and Cumming, 2006; Gu et al., 2008; Smolyanskaya et al., 2015). However, other studies did not find such structure (Jasper et al., 2019; Zhao et al., 2020; Levi et al., 2023), or found substantial unexplained differences across subjects (Goris et al., 2017; Jasper et al., 2019; Lange et al., 2023).

Variability in sensory cortex reflects a complex mixture of decision-related activity (the signal we seek to isolate) and other signals that are not related to perception or the decision-making process per se, but are difficult to control and vary across subjects and tasks. Examples include stimulus expectation (Goris et al., 2017), action planning (Laamerad et al., 2023), feature-based attention (Quinn et al., 2021), and choice- and reward history (Lueckmann et al., 2018; Jasper et al., 2019). So for the moment, we do not know how to create conditions under which an empirical measurement will reliably distinguish the neurons that contribute to perception.

## Acknowledgements

This work was supported by National Institutes of Health Grants EY022428 (to J.A.M. and E.P.S.), EY032102 (to C.M.Z.), and EY032999 (to R.L.T.G).

## Notes

### Competing Interest Statement

The authors have declared no competing interest.

### Summary of Updates

Fixed a formatting issue with original draft

## References

Adams DL, Economides JR, Jocson CM, Horton JC (2007) A biocompatible titanium headpost for stabilizing behaving monkeys. Journal of Neurophysiology 98:993–1001.

Adams DL, Economides JR, Jocson CM, Parker JM, Horton JC (2011) A watertight acrylic-free titanium recording chamber for electrophysiology in behaving monkeys. Journal of Neurophysiology 106:1581–1590.

Alitto HJ, Moore BD, Rathbun DL, Usrey WM (2011) A comparison of visual responses in the lateral geniculate nucleus of alert and anaesthetized macaque monkeys. The Journal of Physiology 589:87–99.

Arcizet F, Jouffrais C, Girard P (2008) Natural textures classification in area V4 of the macaque monkey. Experimental Brain Research 189:109–120.

Bair W, Cavanaugh JR, Movshon JA (2003) Time course and time-distance relationships for surround suppression in macaque V1 neurons. The Journal of Neuroscience: The Official Journal of the Society for Neuroscience 23:7690–7701.

Britten KH, Newsome WT, Shadlen MN, Celebrini S, Movshon JA (1996) A relationship between behavioral choice and the visual responses of neurons in macaque MT. Visual Neuroscience 13:87–100.

Britten KH, Shadlen MN, Newsome WT, Movshon JA (1992) The analysis of visual motion: a comparison of neuronal and psychophysical performance. The Journal of Neuroscience 12:4745–4765.

Cumming BG, Nienborg H (2016) Feedforward and feedback sources of choice probability in neural population responses. Current Opinion in Neurobiology 37:126–132.

Dodd JV, Krug K, Cumming BG, Parker AJ (2001) Perceptually bistable three-dimensional figures evoke high choice probabilities in cortical area MT. The Journal of Neuroscience: The Official Journal of the Society for Neuroscience 21:4809–4821.

Freeman J, Simoncelli EP (2011) Metamers of the ventral stream. Nature Neuroscience 14:1195–1201.

Freeman J, Ziemba CM, Heeger DJ, Simoncelli EP, Movshon JA (2013) A functional and perceptual signature of the second visual area in primates. Nature Neuroscience 16:974–981.

Goris RLT, Simoncelli EP, Movshon JA (2015) Origin and Function of Tuning Diversity in Macaque Visual Cortex. Neuron 88:819–831.

Goris RLT, Ziemba CM, Stine GM, Simoncelli EP, Movshon JA (2017) Dissociation of choice formation and choice-correlated activity in macaque visual cortex. Journal of Neuroscience pp. 3331–16.

Gu Y, Angelaki DE, DeAngelis GC (2008) Neural correlates of multisensory cue integration in macaque MSTd. Nature Neuroscience 11:1201–1210.

Gu Y, DeAngelis GC, Angelaki DE (2007) A functional link between area MSTd and heading perception based on vestibular signals. Nature Neuroscience 10:1038–1047.

Haefner RM, Berkes P, Fiser J (2014) The implications of perception as probabilistic inference for correlated neural variability during behavior. arXiv:1409.0257 [q-bio] 1409.0257.

Haefner RM, Berkes P, Fiser J (2016) Perceptual Decision-Making as Probabilistic Inference by Neural Sampling. Neuron 0.

Hegdé J, Van Essen DC (2004) Temporal dynamics of shape analysis in macaque visual area V2. Journal of Neurophysiology 92:3030–3042.

Hegdé J, Van Essen DC (2007) A comparative study of shape representation in macaque visual areas v2 and v4. Cerebral Cortex (New York, N.Y.: 1991) 17:1100–1116.

Henderson MM, Tarr MJ, Wehbe L (2023) A Texture Statistics Encoding Model Reveals Hierarchical Feature Selectivity across Human Visual Cortex. Journal of Neuroscience 43:4144–4161.

Jasper AI, Tanabe S, Kohn A (2019) Predicting Perceptual Decisions Using Visual Cortical Population Responses and Choice History. Journal of Neuroscience 39:6714–6727.

Jazayeri M, Afraz A (2017) Navigating the Neural Space in Search of the Neural Code. Neuron 93:1003–1014.

Jazayeri M, Wallisch P, Movshon JA (2012) Dynamics of Macaque MT Cell Responses to Grating Triplets. The Journal of Neuroscience 32:8242–8253.

Kohler PJ, Clarke A, Yakovleva A, Liu Y, Norcia AM (2016) Representation of maximally regular textures in human visual cortex. The Journal of Neuroscience 36:714–729.

Kosai Y, El-Shamayleh Y, Fyall AM, Pasupathy A (2014) The role of visual area V4 in the discrimination of partially occluded shapes. The Journal of Neuroscience 34:8570–8584.

Krishna A, Tanabe S, Kohn A (2021) Decision Signals in the Local Field Potentials of Early and Mid-Level Macaque Visual Cortex. Cerebral Cortex 31:169–183.

Laamerad P, Liu LD, Pack CC (2023) Decision-related activity and movement selection in primate visual cortex.

Lange RD, Gómez-Laberge C, Berezovskii VK, Pletenev A, Sherdil A, Hartmann T, Haefner RM, Born RT (2023) Weak evidence for neural correlates of task-switching in macaque V1. Journal of Neurophysiology 129:1021–1044.

Levi AJ, Zhao Y, Park IM, Huk AC (2023) Sensory and Choice Responses in MT Distinct from Motion Encoding. Journal of Neuroscience 43:2090–2103.

Lueckmann JM, Macke JH, Nienborg H (2018) Can Serial Dependencies in Choices and Neural Activity Explain Choice Probabilities? Journal of Neuroscience 38:3495–3506.

Movshon JA, Simoncelli EP (2015) Representation of Naturalistic Image Structure in the Primate Visual Cortex. Cold Spring Harbor Symposia on Quantitative Biology p. 024844.

Newsome WT, Britten KH, Movshon JA (1989) Neuronal correlates of a perceptual decision. Nature 341:52–54.

Nienborg H, Cohen MR, Cumming BG (2012) Decision-related activity in sensory neurons: correlations among neurons and with behavior. Annual Review of Neuroscience 35:463–483.

Nienborg H, Cumming BG (2006) Macaque V2 Neurons, But Not V1 Neurons, Show Choice-Related Activity. The Journal of Neuroscience 26:9567–9578.

Nienborg H, Cumming BG (2009) Decision-related activity in sensory neurons reflects more than a neuron’s causal effect. Nature 459:89–92.

Nienborg H, Cumming BG (2014) Decision-related activity in sensory neurons may depend on the columnar architecture of cerebral cortex. The Journal of Neuroscience: The Official Journal of the Society for Neuroscience 34:3579–3585.

Okazawa G, Tajima S, Komatsu H (2015) Image statistics underlying natural texture selectivity of neurons in macaque V4. Proceedings of the National Academy of Sciences of the United States of America 112:E351–360.

Okazawa G, Tajima S, Komatsu H (2017) Gradual development of visual texture-selective properties between macaque areas V2 and V4. Cerebral Cortex 27:4867–4880.

Parker AJ, Newsome WT (1998) SENSE AND THE SINGLE NEURON: Probing the Physiology of Perception. Annual Review of Neuroscience 21:227–277.

Pitkow X, Liu S, Angelaki DE, DeAngelis GC, Pouget A (2015) How Can Single Sensory Neurons Predict Behavior? Neuron 87:411–423.

Portilla J, Simoncelli EP (2000) A parametric texture model based on joint statistics of complex wavelet coefficients. International Journal of Computer Vision 40:49–70.

Priebe N, Ferster D (2012) Mechanisms of Neuronal Computation in Mammalian Visual Cortex. Neuron 75:194–208.

Prince SJ, Pointon AD, Cumming BG, Parker AJ (2000) The precision of single neuron responses in cortical area V1 during stereoscopic depth judgments. The Journal of Neuroscience: The Official Journal of the Society for Neuroscience 20:3387–3400.

Purushothaman G, Bradley DC (2005) Neural population code for fine perceptual decisions in area MT. Nature Neuroscience 8:99–106.

Quinn KR, Seillier L, Butts DA, Nienborg H (2021) Decision-related feedback in visual cortex lacks spatial selectivity. Nature Communications 12:4473.

Rust NC, Dicarlo JJ (2010) Selectivity and tolerance (“invariance”) both increase as visual information propagates from cortical area V4 to IT. The Journal of Neuroscience: The Official Journal of the Society for Neuroscience 30:12978–12995.

Salzman CD, Britten KH, Newsome WT (1990) Cortical microstimulation influences perceptual judgements of motion direction. Nature 346:174–177.

Shadlen MN, Britten KH, Newsome WT, Movshon JA (1996) A computational analysis of the relationship between neuronal and behavioral responses to visual motion. The Journal of Neuroscience: The Official Journal of the Society for Neuroscience 16:1486–1510.

Shiozaki HM, Tanabe S, Doi T, Fujita I (2012) Neural activity in cortical area V4 underlies fine disparity discrimination. The Journal of Neuroscience 32:3830–3841.

Smith MA, Majaj NJ, Movshon JA (2005) Dynamics of motion signaling by neurons in macaque area MT. Nature Neuroscience 8:220–228.

Smolyanskaya A, Haefner RM, Lomber SG, Born RT (2015) A Modality-Specific Feedforward Component of Choice-Related Activity in MT. Neuron 87:208–219.

Thomas OM, Cumming BG, Parker AJ (2002) A specialization for relative disparity in V2. Nature Neuroscience 5:472–478.

Uka T, DeAngelis GC (2006) Linking Neural Representation to Function in Stereoscopic Depth Perception: Roles of the Middle Temporal Area in Coarse versus Fine Disparity Discrimination. The Journal of Neuroscience 26:6791–6802.

Vacher J, Davila A, Kohn A, Coen-Cagli R (2020) Texture Interpolation for Probing Visual Perception. Advances in Neural Information Processing Systems 33:22146–22157.

Wimmer K, Compte A, Roxin A, Peixoto D, Renart A, de la Rocha J (2015) Sensory integration dynamics in a hierarchical network explains choice probabilities in cortical area MT. Nature Communications 6.

Yu X, Gu Y (2018) Probing Sensory Readout via Combined Choice-Correlation Measures and Microstimulation Perturbation. Neuron 100:715–727.e5.

Zaidel A, DeAngelis GC, Angelaki DE (2017) Decoupled choice-driven and stimulus-related activity in parietal neurons may be misrepresented by choice probabilities. Nature Communications 8:715.

Zhao Y, Yates JL, Levi AJ, Huk AC, Park IM (2020) Stimulus-choice (mis)alignment in primate area MT. PLOS Computational Biology 16:e1007614.

Ziemba CM, Freeman J, Movshon JA, Simoncelli EP (2016) Selectivity and tolerance for visual texture in macaque V2. Proceedings of the National Academy of Sciences 113:E3140–E3149.

Ziemba CM, Freeman J, Simoncelli EP, Movshon JA (2018) Contextual modulation of sensitivity to naturalistic image structure in macaque V2. Journal of Neurophysiology 120:409–420.

Ziemba CM, Perez RK, Pai J, Kelly JG, Hallum LE, Shooner C, Movshon JA (2019) Laminar differences in responses to naturalistic texture in macaque V1 and V2. Journal of Neuroscience 39:9748–9756.

Ziemba CM, Simoncelli EP (2021) Opposing effects of selectivity and invariance in peripheral vision. Nature Communications 12:4597.

